# Dynamic changes in mRNA nucleocytoplasmic localization in the nitrate response of Arabidopsis roots

**DOI:** 10.1101/2022.01.07.475360

**Authors:** Alejandro Fonseca, Tomás C. Moyano, Stefanie Rosa, Rodrigo A. Gutiérrez

## Abstract

Nitrate (NO_3_^-^) is a signaling molecule that regulates gene expression in plants. The nitrate response has been extensively characterized at the transcriptome level. However, we know little about RNA nucleocytoplasmic dynamics during nitrate response. To understand the role of mRNA localization during the nitrate response, we isolated mRNA from the nucleus, cytoplasm, and whole-cells from nitrate-treated Arabidopsis roots and performed RNA-seq. We identified 402 differentially localized transcripts (DLTs) in response to nitrate. DLTs were enriched in GO-terms related to metabolism, response to stimulus, and transport. DLTs showed five localization patterns: nuclear reduction, cytoplasmic reduction, nuclear accumulation, cytoplasmic accumulation, or delayed-cytoplasmic accumulation in response to nitrate. DLTs exhibited large changes in RNA polymerase II occupancy of cognate genes and high mRNA turnover rates, indicating these are rapidly replaced mRNAs. The *NITRATE REDUCTASE 1* (*NIA1*) transcript exhibited the largest changes in synthesis and decay. Using single-molecule RNA FISH, we showed that *NIA1* nuclear accumulation occurs mainly at transcription sites. The decay profiles for *NIA1* showed a higher half-life when the transcript accumulated in the nucleus than in the cytoplasm. We propose that regulating nucleocytoplasmic mRNA distribution allows tuning transcript availability of fastly replaced mRNAs, controlling plants’ adaptive response to nitrogen nutrient signals.

## INTRODUCTION

Nitrogen (N) is an essential macronutrient whose availability limits growth and development in plants (Andrews et al., 2013; Gutiérrez, 2013; Fredes et al., 2019; Araus et al., 2020; Vidal et al., 2020; Alvarez et al., 2021). Nitrate is the most abundant source of N in agricultural soils (Owen and Jones, 2001). Nitrate acts as a signaling molecule (Scheible et al., 1997; Wang et al., 2004) that initiates a signal transduction cascade (Undurraga et al., 2017; Vidal et al., 2020). The dual-affinity transceptor NPF6.3/NRT1.1 senses nitrate (Ho et al., 2009). Different regulatory factors, at the local and systemic level, orchestrate downstream responses affecting nutrient metabolism and a series of developmental processes associated with root development (Forde and Walch-liu, 2009; Vidal et al., 2010; Gruber et al., 2013; Alvarez et al., 2014; O’Brien et al., 2016; Canales et al., 2017), shoot development (Rahayu et al., 2005; Landrein et al., 2018; Poitout et al., 2018; Moreno et al., 2020), seed dormancy (Alboresi et al., 2005; Yan et al., 2016), and flowering time (Castro Marín et al., 2011; Gras et al., 2018). In addition to the NRT1.1 transceptor, critical components in the nitrate signaling pathway include CIPK23 kinase (Liu and Tsay, 2003), calcium as a second messenger (Riveras et al., 2015), and a myriad of transcription factors controlling transcriptional responses such as NLP7 (Marchive et al., 2013; Alvarez et al., 2020), TGA1 and TGA4 (Alvarez et al., 2014; Swift et al., 2020), NAC4 (Vidal et al., 2013b), SPL9 (Krouk et al., 2010), HRS1 and HHO1 (Medici et al., 2015; Maeda et al., 2018), NRG2 (Xu et al., 2016), TCP20 (Guan et al., 2017), and CRF4 (Varala et al., 2018).

In eukaryotic cells, mRNA synthesis and processing occur in the nucleus, and translation mainly in the cytoplasm (Martin and Koonin, 2006). This compartmentalization of mRNA processes allows for sophisticated regulation of gene expression (Wickramasinghe and Laskey, 2015). The nucleocytoplasmic dynamic of transcripts is mainly determined by synthesis, export, and decay factors (Bahar Halpern et al., 2015; Hansen et al., 2018). Synthesis and decay rates have been quantified at the genome-wide level in yeast (Miller et al., 2011; Sun et al., 2012; Eser et al., 2014), mouse (Schwanhausser et al., 2011; Tippmann et al., 2012; Rabani et al., 2014; Jovanovic et al., 2015), flies (Chen and Van Steensel, 2017), and plants (Gutierrez et al., 2002; Sorenson et al., 2018; Szabo et al., 2020). These results indicate that synthesis and decay rates contribute to mRNA steady-state levels in a species-specific manner. The sequencing of RNA from cellular fractions of different eukaryotic species showed that transcripts are asymmetrically distributed between the nucleus and cytoplasm (Barthelson et al., 2007; Djebali et al., 2012; Solnestam et al., 2012; Bahar Halpern et al., 2015; Battich et al., 2015; Chen and Van Steensel, 2017; Kim et al., 2017; Pastro et al., 2017; Abdelmoez et al., 2018; Benoit Bouvrette et al., 2018; Lee and Bailey-Serres, 2019; Palovaara and Weijers, 2019; Reynoso et al., 2019). Controlling mRNA nuclear export to change the availability of transcripts for translation allows the cell to fine-tune gene expression according to environmental and cellular requirements (Parry, 2015; Wickramasinghe and Laskey, 2015; Chen and Van Steensel, 2017; Yang et al., 2017; Lee and Bailey-Serres, 2019).

In plants, the export-machinery components are more diverse than in yeast or animals (Yelina et al., 2010; Pfaff et al., 2018), suggesting their ability to regulate cytoplasmic mRNA levels in response to a stimulus is more versatile (Ehrnsberger and Grasser, 2019). Some studies have shown that subsets of mRNAs display particular nucleocytoplasmic distributions during different plant processes, such as cell cycle control (Yang et al., 2017), ethylene signaling (Chen et al., 2019), RNA-directed DNA methylation (Choudury et al., 2019), and stress response (Yeap et al., 2019). However, the mRNA nucleocytoplasmic dynamics at the genome-wide level have only been described in response to flooding stress (Lee and Bailey-Serres, 2019; Reynoso et al., 2019). Genome-wide changes in gene expression in response to nitrate treatments have been thoroughly characterized in several studies (Wang et al., 2003; Wang et al., 2004; Gutiérrez et al., 2007; Wang et al., 2007; Gifford et al., 2008; Hu et al., 2009; Krouk et al., 2009; Krouk et al., 2010; Patterson et al., 2010; Ruffel et al., 2011; Vidal et al., 2013a; Alvarez et al., 2014; Walker et al., 2017; Gaudinier et al., 2018; Varala et al., 2018; Alvarez et al., 2019; Moreno et al., 2020; Swift et al., 2020). However, we currently lack an understanding of the importance of mRNA nucleocytoplasmic dynamics in the nitrate response.

In this work, we aimed to understand the nucleocytoplasmic dynamics of mRNAs in response to nitrate treatments. We used RNA-seq analysis from nuclear, cytoplasmic, and total fractions to identify differentially localized transcripts (DLTs) in response to nitrate treatment. Integrated analysis of our genome-wide data showed that DLTs have significant synthesis and decay rate changes, indicating these mRNAs are rapidly replaced during the nitrate response. Our results propose a role for mRNA nuclear export in regulating gene expression, which is critical for the plants’ ability to adapt to nutritional changes

## RESULTS

### Identification of differentially expressed genes in response to nitrate in subcellular fractions

To understand the distribution of mRNAs between nucleus and cytoplasm, we analyzed mRNA levels in response to nitrate treatments in nuclear and cytoplasmic subcellular fractions. Total RNA was obtained from nuclear, cytoplasmic, and total fractions. RNA samples were prepared from Arabidopsis roots 0, 20, 60, and 120 minutes after nitrate or control treatments. We quantified RNA levels for selected transcripts using RT-qPCR as a control experiment. As shown in Supplemental Figure 1, we observed enrichment of unprocessed transcripts in the nuclear fraction and a significant reduction in the cytoplasmic fraction compared to total RNA. We performed RNA-seq analysis with the material obtained from subcellular and total fractions. We analyzed three independent replicates for each condition (separate plant material grown independently). Supplemental Table 1 summarizes quality parameters for all libraries. We found high reproducibility among replicate experiments with a mean Pearson correlation of 0.985 ± 0.003 (Supplemental Table 1). Sequence data were filtered by quality, mapped to the Araport11 Arabidopsis genome, and normalized as detailed in Materials and Methods.

To identify genes with changes in their mRNA levels in response to the treatments, we performed analyses of variance (ANOVA) for the total or each subcellular fraction separately. Our ANOVA model evaluated the effect of treatments (KCl and KNO_3_), time (20, 60, 120 min), or their interactions (Supplemental Figure 2A-C). We selected significant models with a p-value < 0.01 after FDR correction. We found 6,006 genes whose mRNA levels changed by the treatment or interactions in the total fraction (Supplemental Data Set 1). Analysis of gene ontology (GO) terms identified over-represented biological processes in response to the nitrate treatment, such as nitrate response, nitrate transport, nitrate assimilation, development, response to hormones, amino acid metabolism, nucleotide metabolism, carbon metabolism, among others (Supplemental Data Set 2). We identified 4,445 differentially expressed genes in the nuclear or cytoplasmic fractions (Supplemental Figure 2B-C, Supplemental Data Set 1), where 1,183 genes did not show significant changes in the total fraction (Supplemental Figure 3). This result indicates that the nitrate-response analysis from subcellular fractions provides complementary information to the analysis of total RNA, detecting genes whose mRNAs accumulate specifically in one fraction and cannot be easily detected in total RNA. Interestingly, despite extensive transcriptome analysis of the nitrate response in Arabidopsis, 445 genes differentially expressed in this study have not been identified previously (Supplemental Figure 4). These genes code for proteins involved in growth and development (e.g., *AUXIN RESISTANT 1*, *GROWTH-REGULATING FACTOR 2* and *BRASSINOSTEROID-INSENSITIVE 2*), cell cycle (e.g*., INCREASED LEVEL OF POLYPLOIDY1-1D*), signaling (e.g., *CBL-INTERACTING PROTEIN KINASE 19* and *MAP KINASE 7*), protein modification (e.g., *SUMO-ACTIVATING ENZYME 2* and *UBIQUITIN PROTEIN LIGASE 6*), nitrogen compound metabolism (e.g., *METHIONINE OVER-ACCUMULATOR 2* and *NICOTINAMIDASE 1*), response to stress (e.g., *ANKYRIN REPEAT-CONTAINING PROTEIN 2* and *C-REPEAT/DRE BINDING FACTOR 1*), among other functions. Furthermore, uncharacterized long non-coding RNAs (*AT1G06103, AT1G08697, AT2G09525, AT3G05055, AT4G06085, AT4G06935, AT4G06945, AT5G06585, AT5G09125*) and antisense long non-coding RNAs (*AT1G34844, AT1G67328, AT2G07275, AT3G01205, AT3G09575, AT4G05015, AT4G22233, AT5G01375, AT5G08235, AT5G09595*) were also regulated in response to nitrate in the subcellular fractions (Supplemental Data Set 1). These genes represent new components of the nitrate response and could contribute to the plant adaptation to N availability changes.

### Differentially localized transcripts in response to nitrate treatments

We calculated the delta between normalized counts in nuclear and cytoplasmic fractions (ΔNC) to identify genes that change their nucleocytoplasmic distribution in response to nitrate. These ΔNC values were used for two-way ANOVA analysis to evaluate the effect of the treatment, time, or their interactions in mRNA nucleocytoplasmic distribution. We selected significant models with a p-value < 0.01 after FDR correction. <u>D</u>ifferentially localized transcripts (DLTs) in response to nitrate were defined as transcripts whose ΔNC values depend on the treatment or the treatment-time interactions (p-value<0.01). We identified 402 DLTs in response to nitrate treatments in Arabidopsis roots (Supplemental Figure 2D, Supplemental Data Set 3). mRNA levels for DLTs in the total fraction should be a combination of the levels we measured in each fraction. To confirm this assumption, we estimated a ’reconstituted cell’ count for each DLT by simply adding nuclear and cytoplasmic normalized counts (Figure 1A). A high correlation (Pearson correlation value of 0.99) was observed when ’reconstituted cell’ counts (i.e., Nuclear+Cytoplasmic levels) were compared with mRNA levels obtained in the total fraction, validating our experimental approach and data analysis procedure (Figure 1B). Twenty-two percent (88/402) of DLTs were not identified as regulated in the total RNA fraction (Supplemental Figure 5). Some of these genes have biological functions in transcriptional regulation (e.g., *ERF1*, *ERF105*, and *AFP3*), nutrient metabolism (e.g., *CYANASE* – *CYN -* and *SERINE ACETYLTRANSFERASE 2;1* - *SERAT2;1*), auxin response (*SAUR-59*), and root development (e.g., *POPCORN*) (Supplemental Data Set 3).

**Figure 1.**
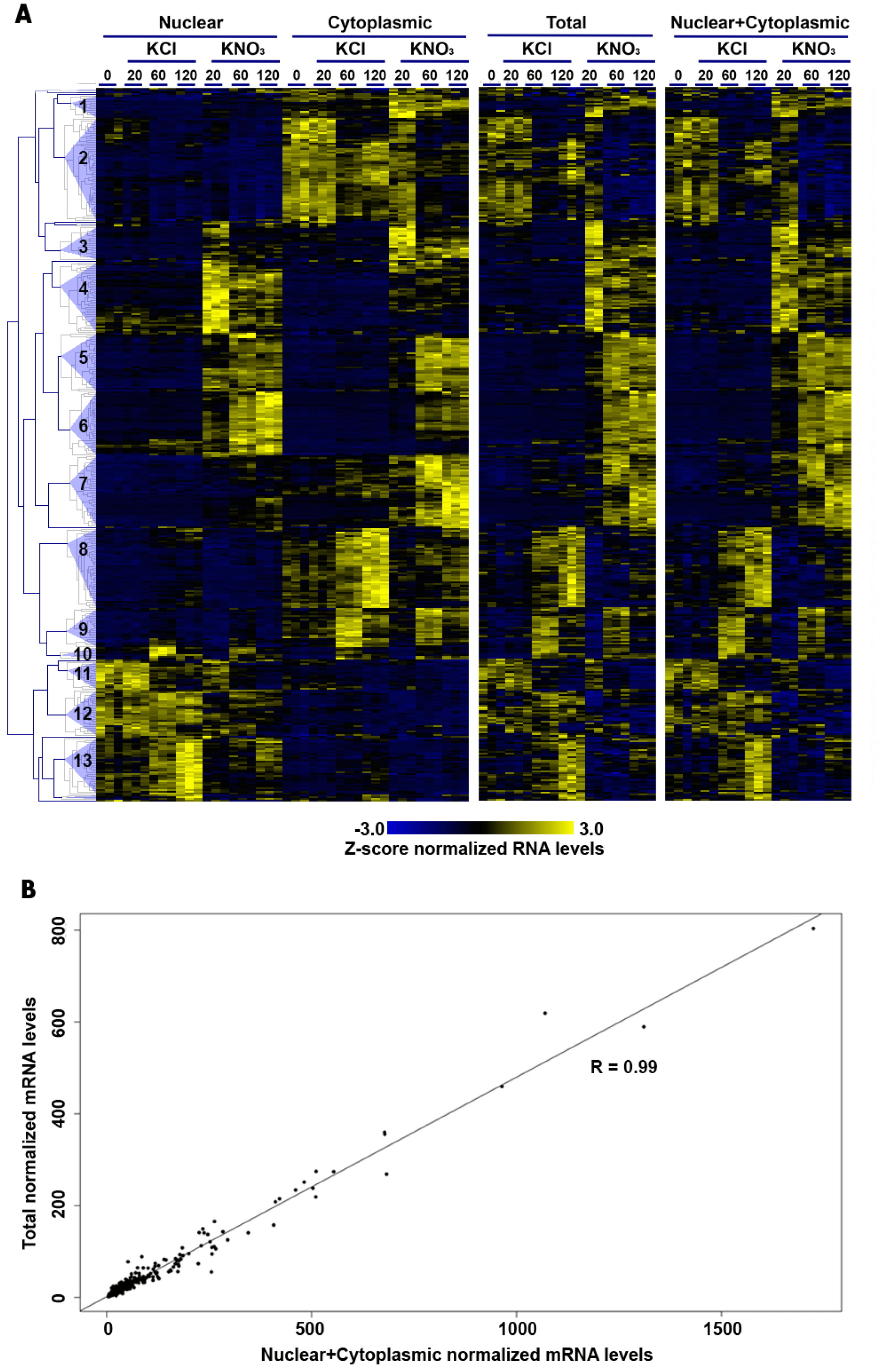
Differentially localized transcripts (DLTs) in response to nitrate treatments. Heatmap with mRNA levels in cellular fractions for DLTs in response to nitrate treatments. Genes were hierarchically clustered using the correlation of mRNA levels in the nuclear and cytoplasmic fractions (panel I). mRNA levels in the total fraction are shown in panel II. We calculated Nuclear+Cytoplasmic mRNA levels using data from each fraction, which are shown in panel III. Cluster numbers are indicated on the dendrogram to the left of the heatmap. Each column represents the mRNA levels for one replicate under each condition. **(B)** Scatter plot for comparing the mean mRNA levels of DLTs in the Total and Nuclear+Cytoplasmic data. The Pearson correlation (R) is indicated.

To understand the nitrate response dynamics of mRNA levels in the different fractions, we performed a hierarchical clustering analysis for the 402 DLTs. We obtained 13 clusters with five or more genes, including 389 DLTs (Figure 1A). As shown in Figure 2 and Supplemental Figure 6, these 13 clusters correspond to five different localization patterns in response to the nitrate treatment: Nuclear reduction (NR), containing 81 genes with decreasing RNA levels in the nucleus; Cytoplasmic reduction (CR), with 125 genes with decreasing RNA levels in the cytoplasm; Nuclear accumulation (NA), containing 76 genes with increasing levels in the nucleus; Cytoplasmic accumulation (CA), with 72 genes with increasing levels in the cytoplasm; and Delayed-cytoplasmic accumulation (D), containing 33 genes which showed nuclear enrichment at 20 min of treatment and cytoplasmic enrichment at later times (Figure 2, Supplemental Data Set 3).

**Figure 2.**
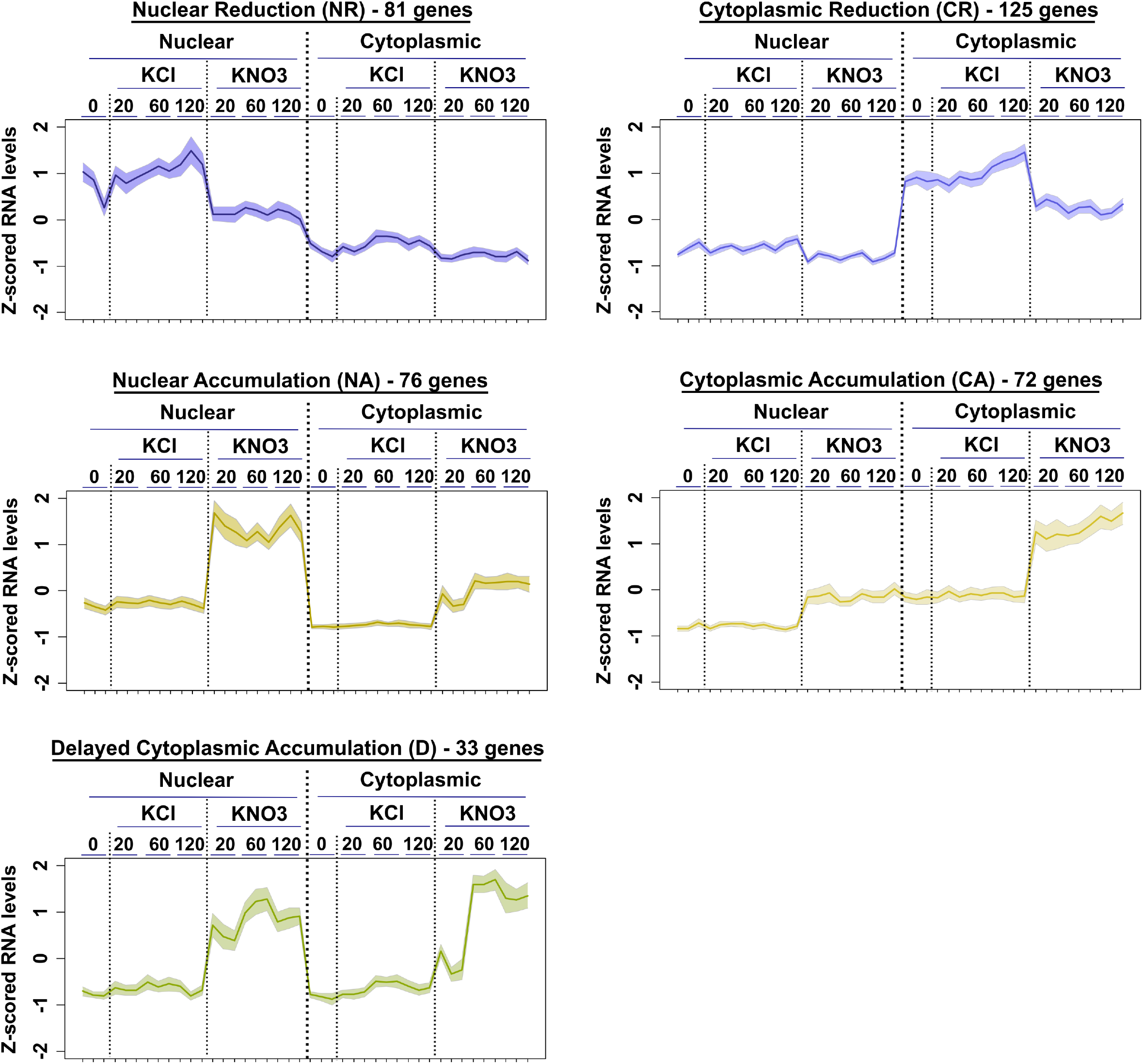
DLT localization patterns in response to nitrate treatments. Five different localization patterns in response to nitrate treatments. NR – Nuclear reduction, CR – Cytoplasmic reduction, NA – Nuclear accumulation, CA – Cytoplasmic accumulation, and D – Delayed-cytoplasmic accumulation. DLTs in response to nitrate treatments were separated by the cellular fraction where the main changes were observed (Nuclear or cytoplasmic) and if they showed an accumulation or reduction of mRNA levels in the nitrate condition. Graphs show mean values of z-scored normalized mRNA levels (line) and 95% confidence interval for mean values of each DLT for the three independent experiments (shadow).

Significantly over-represented gene ontology (GO) and Kyoto Encyclopedia of Genes and Genome (KEGG) terms in DLTs include metabolic processes (cofactor, nitrogen compound, carbohydrate, glycerolipid, energy metabolism), localization (anion, amine, and organic acid transport), and response to stimulus functions (Figure 3, Supplemental Data Set 4, Supplemental Data Set 5). We identified anion transport, histidine biosynthesis, and nucleotide biosynthesis in the nuclear accumulation pattern. In the cytoplasmic accumulation pattern, we found nicotianamine biosynthesis, regulation of organic acid and amino acid export, and sulfur metabolism pathways over-represented. Besides, we found that the following biological processes were over-represented among DLTs with a delayed-cytoplasmic accumulation pattern: carbohydrate metabolism (specifically glycolysis/gluconeogenesis and TCA cycle), nitrogen compound metabolism, cofactor metabolic process, and cellular amino acid biosynthesis. In the cytoplasmic reduction pattern, we found the response to stimulus and glycerolipid metabolism functions. We did not observe over-represented terms for the nuclear reduction pattern (Figure 3, Supplemental Data Set 4, Supplemental Data Set 5). These results demonstrate that mRNAs with relevant functions for the nitrate response are differentially distributed in the cellular fractions.

**Figure 3.**
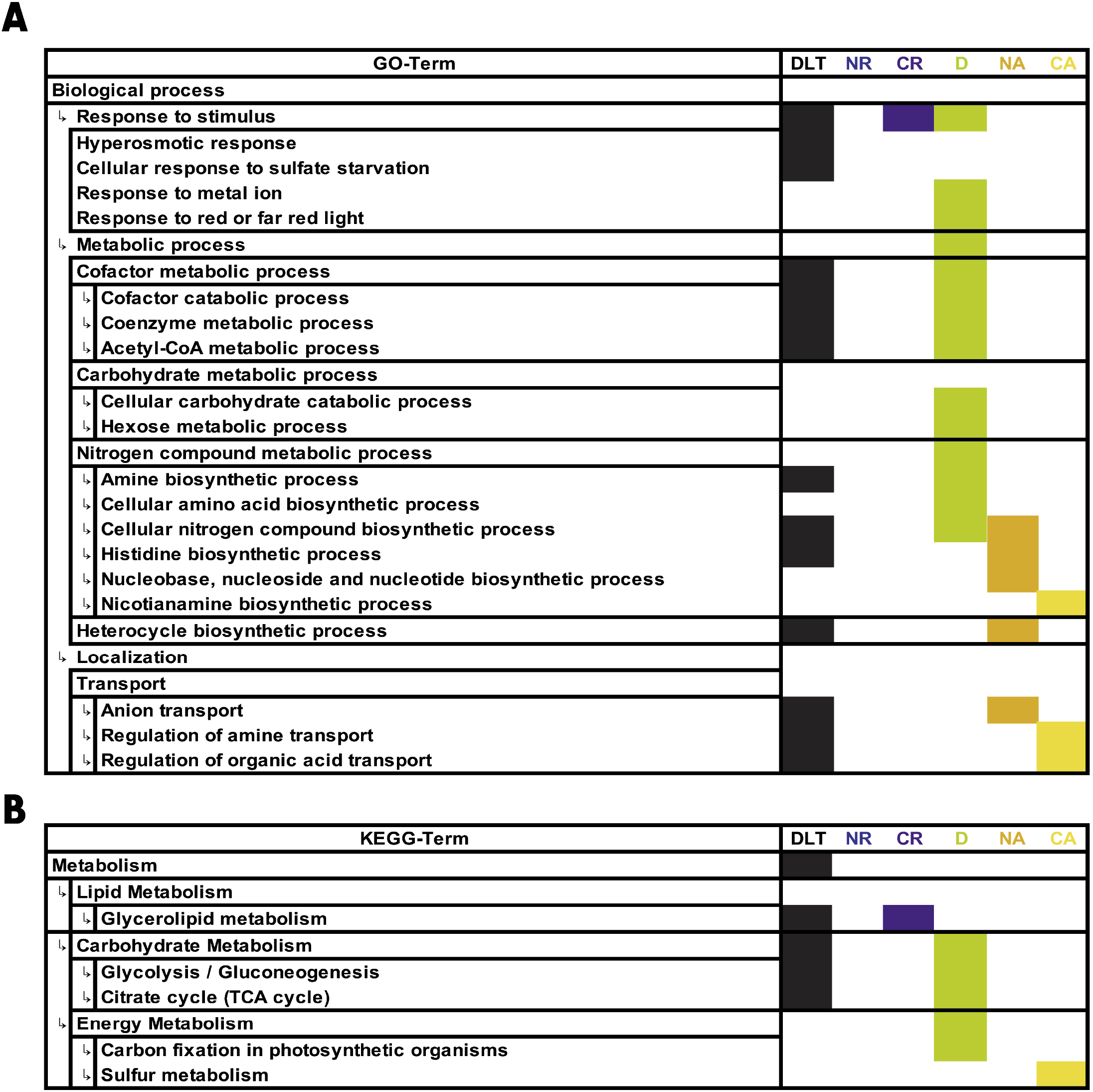
Over-represented terms in DLTs in response to nitrate treatments. Summary of significant (p<0.05) over-representation of **(A)** GO, and **(B)** KEGG-Terms enriched in the lists of all DLTs, or the DLT localization patterns (NR – Nuclear reduction, CR – Cytoplasmic reduction, NA – Nuclear accumulation, D – Delayed-cytoplasmic accumulation, and CA – Cytoplasmic accumulation) according to VirtualPlant output (Katari et al., 2010). GO-terms were summarized by non-redundant 5 and 6 levels using REVIGO (Supek et al., 2011).

We selected two representative genes from each localization pattern to validate the RNA-seq data. We measured mRNA levels by RT-qPCR in the time-point where the most significant differences were observed between cellular fractions (Supplemental Figure 7). The selected genes were: *MPK9* and *SDR2* for the nuclear reduction pattern; *SUFE2* and *RCAR1* for the cytoplasmic reduction pattern; *NRT2.2* and *BCA4* for the nuclear accumulation pattern; *BZIP3* and *AT1G49230* for the cytoplasmic accumulation pattern; and *NIA1* and *IDH1* for the delayed-cytoplasmic accumulation pattern. We validated the differential mRNA localization pattern in all cases (Supplemental Figure 7), confirming the RNA-seq and data analysis results.

Our cell-fractionation/RNA-seq strategy allowed us to identify transcripts with differential localization in the nucleus and cytoplasm in response to nitrate. The corresponding genes have relevant functions for the nitrate response, suggesting that controlling nucleocytoplasmic localization is an important mechanism to regulate gene expression in response to nitrate. Moreover, more than 20% of these genes have not been previously characterized in the plant’s N nutrients response.

### Transcripts from DLT localization patterns showed characteristic structural features

To evaluate whether specific sequence features could be associated with the differential mRNA localization, we evaluated features described to associate with nucleocytoplasmic levels in other species (Palazzo and Lee, 2018). Sequence features could be related to RNA binding protein recognition leading to different RNA processes (Chen and Van Steensel, 2017; Benoit Bouvrette et al., 2018; Dedow and Bailey-Serres, 2019). Furthermore, nuclear retention is associated with RNA length and splicing events (Monteuuis et al., 2019; Mordstein et al., 2020). We found differences in length, guanine-cytosine (GC) content, and splicing junction density in DLTs as compared to transcripts without differences in the nucleocytoplasmic distribution in response to nitrate treatment (Figure 4 and Supplemental Figures 8 and 9). Cytoplasmic accumulated transcripts showed shorter RNAs and lower GC content in their exonic regions than transcripts induced in response to nitrate but are not differentially localized (Figure 4A-B). These differences are mainly due to shorter CDS regions and lower GC content in the UTRs (Supplemental Figure 8A-F). Cytoplasmic reduced transcripts also showed shorter exonic regions than transcripts repressed in response to nitrate but are not differentially localized (Figure 4A-B, Supplemental Figure 8A-B). To evaluate whether these sequence features could be associated with RNA secondary structure formation differences, we predicted RNA folding energy *in silico* using the RNAfold software. DLTs with cytoplasmic accumulation or reduction patterns exhibited less stable mRNA structures than RNAs that respond to the treatment in the total fraction (induction or repression, respectively) (Supplemental Figure 8G). These differences are also observed in the cytoplasmic accumulation pattern when only the UTRs sequences were analyzed (Supplemental Figure 8H-I). Besides, we observed differences in splicing junction density in DLTs as compared to transcripts without differences in nucleocytoplasmic localization in response to nitrate treatments (Figure 4C). Transcripts in the cytoplasmic reduction pattern showed lower splicing junction density than repressed transcripts in the total fraction. Moreover, transcripts in the nuclear accumulation and delayed-cytoplasmic accumulation patterns also showed higher splicing junction density as compared to induced transcripts in the total fraction. On the other hand, transcripts in the cytoplasmic accumulation pattern showed the lowest splicing junction density values among DLTs (Figure 4C).

**Figure 4.**
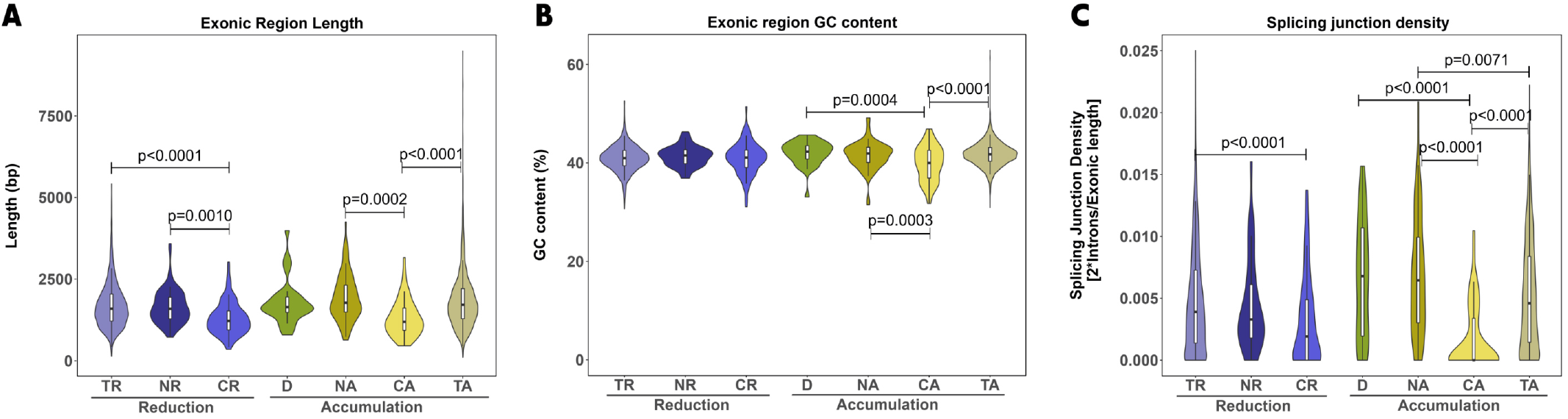
Differences in sequence-related features of DLTs in response to nitrate treatments. Sequence features of the most abundant isoform for nitrate-regulated genes, according to Araport11 annotation. Violin plots show the distribution of **(A)** exonic region length, **(B)** exonic region GC content, and **(C)** splicing junction density for DLTs in each pattern (NR – Nuclear reduction, CR – Cytoplasmic reduction, NA – Nuclear accumulation, D – Delayed-cytoplasmic accumulation, and CA – Cytoplasmic accumulation). We also include nitrate-regulated genes in the total fraction that are not differentially localized as a control (TA – Total accumulated, or TR – Total reduced). Boxes inside show the interquartile range (IQR – 25-75%), the horizontal line indicates the median value. Whiskers show the ±1.58xIQR value. We compared the distributions using one-way ANOVA and Dunnett post-test. We include p-values and brackets to highlight relevant comparisons.

These results show that transcripts with differential localization in response to nitrate have characteristic sequence features. These sequence features have been associated with the modulation of nucleocytoplasmic distribution in yeast and animal systems (Palazzo and Lee, 2018). Our results suggest similar mechanisms may be implicated in the differential localization of plant transcripts.

### Increased RNA polymerase II occupancy is associated with induced DLT genes

Synthesis is one of the most critical processes determining nucleocytoplasmic mRNA levels inside the cell (Bahar Halpern et al., 2015; Hansen et al., 2018). Therefore, we mined published data from our group obtained under the same experimental conditions (Alvarez et al., 2019) to evaluate whether DLTs exhibit specific synthesis changes during nitrate treatment (Figure 5A). We analyzed changes in the RNPII occupancy 12 min after nitrate treatments (Figure 5A, Supplemental Data Set 6). Most of the repressed genes in response to the nitrate did not exhibit changes in RNPII occupancy. We did not observe differences between transcripts with nuclear reduction or cytoplasmic reduction as compared to repressed genes in the total fraction. On the contrary, induced genes by the nitrate treatments also exhibited increased RNPII occupancy. Transcripts with nuclear, cytoplasmic, and delayed-cytoplasmic accumulation showed higher values than transcripts of induced genes that are not differentially localized in response to nitrate.

**Figure 5.**
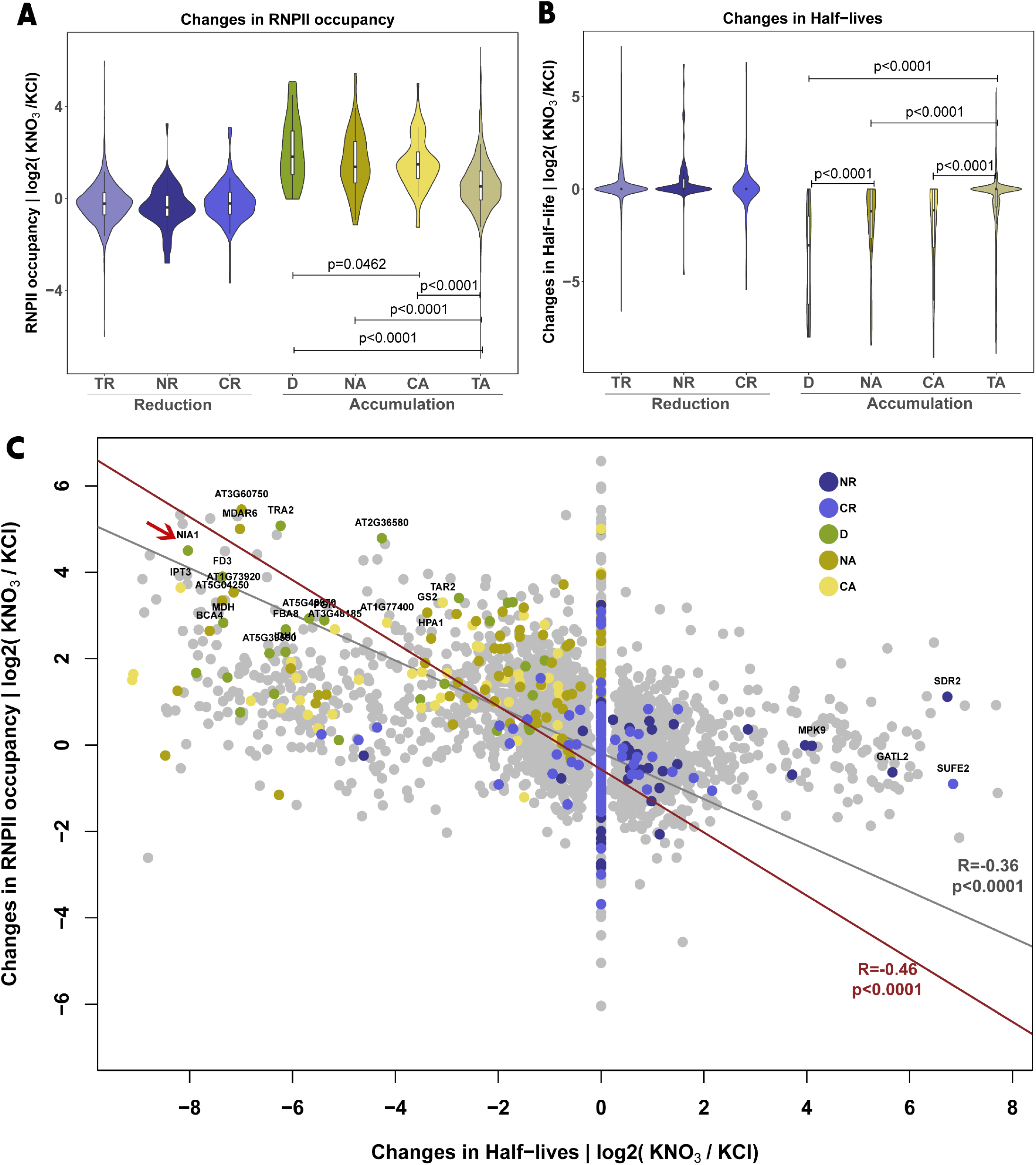
Changes in RNA polymerase II (RNPII) occupancy and half-lives for DLTs in response to nitrate treatments. **(A)** Changes in RNPII occupancy after 12 min and **(B)** Changes in half-lives after 120 min of nitrate treatment. Violin plots show the distribution for DLTs in each pattern (NR – Nuclear reduction, CR – Cytoplasmic reduction, NA – Nuclear accumulation, D – Delayed-cytoplasmic accumulation, and CA – Cytoplasmic accumulation). We also include nitrate-regulated genes in the total fraction that are not differentially localized as a control (TA – Total accumulated, or TR – Total reduced). Boxes inside show the interquartile range (IQR – 25-75%), indicating the median value as a horizontal line. Whiskers show the ±1.58xIQR value. We compared the distributions using one-way ANOVA and Dunnett post-test. We include p-values and brackets to highlight relevant comparisons. **(C)** Scatter plot showing the relationship between changes in RNPII occupancy and half-lives for all genes that respond to nitrate in any cellular fraction. Linear regression and Pearson correlation coefficient are indicated for all data (gray) and DLTs only (red). The red arrow shows *NITRATE REDUCTASE 1* (*NIA1*) as the DLT with the biggest changes in RNPII occupancy as well as half-life values.

These results indicate that an increase in mRNA synthesis rate leads to nuclear accumulation. Interestingly, DLTs with cytoplasmic accumulation, which also exhibited a significant increment in RNPII occupancy, require other regulatory mechanisms (e.g., increased nuclear-to-cytoplasmic transport) to explain their nucleocytoplasmic distribution.

### Negative correlation between mRNA decay rates and mRNA accumulation for DLTs

In addition to synthesis, decay is also essential to determine nucleocytoplasmic mRNA distribution (Bahar Halpern et al., 2015; Hansen et al., 2018). We measured global mRNA decay rates and estimated half-lives using RNA-seq of rRNA-depleted samples. We extracted total RNA from nitrate or control-treated roots in the presence of cordycepin as done previously (Gutierrez et al., 2002; Nagarajan et al., 2019). Sequence data were filtered by quality, mapped to the Araport11 Arabidopsis genome, and counts were normalized to analyze mRNA decay profiles (Materials and Methods) (Supplemental Figure 10). Normalized counts were used for modeling decay rates utilizing an exponential adjustment for RNA levels as a function of time (Materials and Methods). Figure 5B and Supplemental Data Set 7 show changes in mRNA half-lives for each DLT pattern and nitrate-responsive transcripts not differentially localized in the total fraction. Most repressed mRNAs did not change half-lives in response to the treatments. In contrast, transcripts in the delayed-cytoplasmic accumulation, nuclear accumulation, and cytoplasmic accumulation patterns showed significantly faster turnover rates in response to the nitrate treatments (Figure 5B, Supplemental Data Set 7). Moreover, transcripts with delayed-cytoplasmic accumulation showed significantly higher destabilization than those from the nuclear accumulation pattern, indicating the cytoplasmic accumulation leads to faster turnover rates of these transcripts in response to the nitrate treatment.

Interestingly, we found a significant negative correlation when we compared changes in RNPII occupancy and half-lives (Figure 5C) for all nitrate-responsive genes (Pearson correlation = -0.36, p <0.0001). We found an even stronger negative correlation when only DLTs were included in the comparison (Pearson correlation = -0.48, p<0.0001). We calculated the mean rank for RNPII occupancy and half-live changes and found that the top 5% were primarily DLTs in the delayed-cytoplasmic accumulation pattern (green dots in the top left quadrant Figure 5C Supplemental Data Set 8). Among these, the mRNA that encodes the nitrate reduction enzyme NIA1 stood out as the transcript with the biggest differences (red arrow in Figure 5C).

These results reinforce that synthesis, decay rates, and nucleocytoplasmic distribution of DLTs in response to nitrate are connected processes. Furthermore, the negative correlation between RNPII occupancy and half-life changes indicates that induced DLTs are molecules with a rapid replacement in response to the nitrate treatment. This result suggests a role for nucleocytoplasmic dynamics in controlling gene expression, especially for those with delayed-cytoplasmic accumulation patterns (e.g., *NIA1*).

### *NIA1* delayed-cytoplasmic accumulation allows extending the mRNA half-life after a strong transcriptional activation

We selected the *NIA1* transcript for validation and further characterization of extreme DLT patterns to obtain insights into the role of nucleocytoplasmic dynamics during the nitrate treatment. We selected this transcript for three main reasons: (1) the importance of the *NIA1* gene for the nitrate response, (2) the nitrate-induced changes in mRNA synthesis and decay described in the previous section, and (3) its delayed-cytoplasmic accumulation, which allow us to study its nuclear (20 min of treatment) and then its cytoplasmic (60-120 min of treatment) accumulation phases (Figure 6A). *NIA1* mRNA localization at the subcellular level was evaluated by single-molecule RNA FISH (smFISH) in root tip cells (Figure 6B). The number of nuclear, cytoplasmic, and total mRNA molecules was calculated using the FISHquant software (Material and Methods). As shown in Figure 6B, the probe signal showed two different patterns: (1) small fluorescent dots corresponding to nucleoplasmic or cytoplasmic single-molecules (2) big fluorescent foci located in the nucleus, which correspond to active transcription sites. These big nuclear foci disappeared after cordycepin treatments, confirming they are associated with active transcription (Supplemental Figure 11A). We calculated the number of transcripts for whole cells considering single-molecule counts and the estimated number of molecules in transcription sites. As expected, we observed a higher number of RNA molecules per cell area in the nitrate than the control-treated cells (Figure 6C). We found that the number of molecules increases more in the cytoplasm than the nucleus at 120 min in the nitrate condition (Figure 6D-E), which is consistent with the delayed-cytoplasmic accumulation DLT pattern described for this transcript. This result validates the differential localization pattern for *NIA1* mRNA observed in the RNA-seq data at a cellular level. Higher RNA levels at 20 minutes of nitrate treatment are observed in the transcription sites but not in the nucleoplasm (Figure 6F-G). This result indicates that differences in mRNA nuclear levels for *NIA1* derive from more nascent RNAs or mRNAs accumulating at transcription sites. This phenomenon can be explained for the activation of more synthesis loci, observing a higher number of active transcription sites per cell at 20 min as compared to the 120 min after nitrate treatments (Supplemental Figure 11B), more than differences in the intensity of these loci (Supplemental Figure 11C). These results indicate that early nuclear accumulation of *NIA1* is associated with a strong transcriptional activation. After some minutes, RNA accumulation at transcription sites diminishes, and mRNAs accumulate in the cytoplasm, presumably due to increased release from the DNA locus and nuclear export rates.

**Figure 6.**
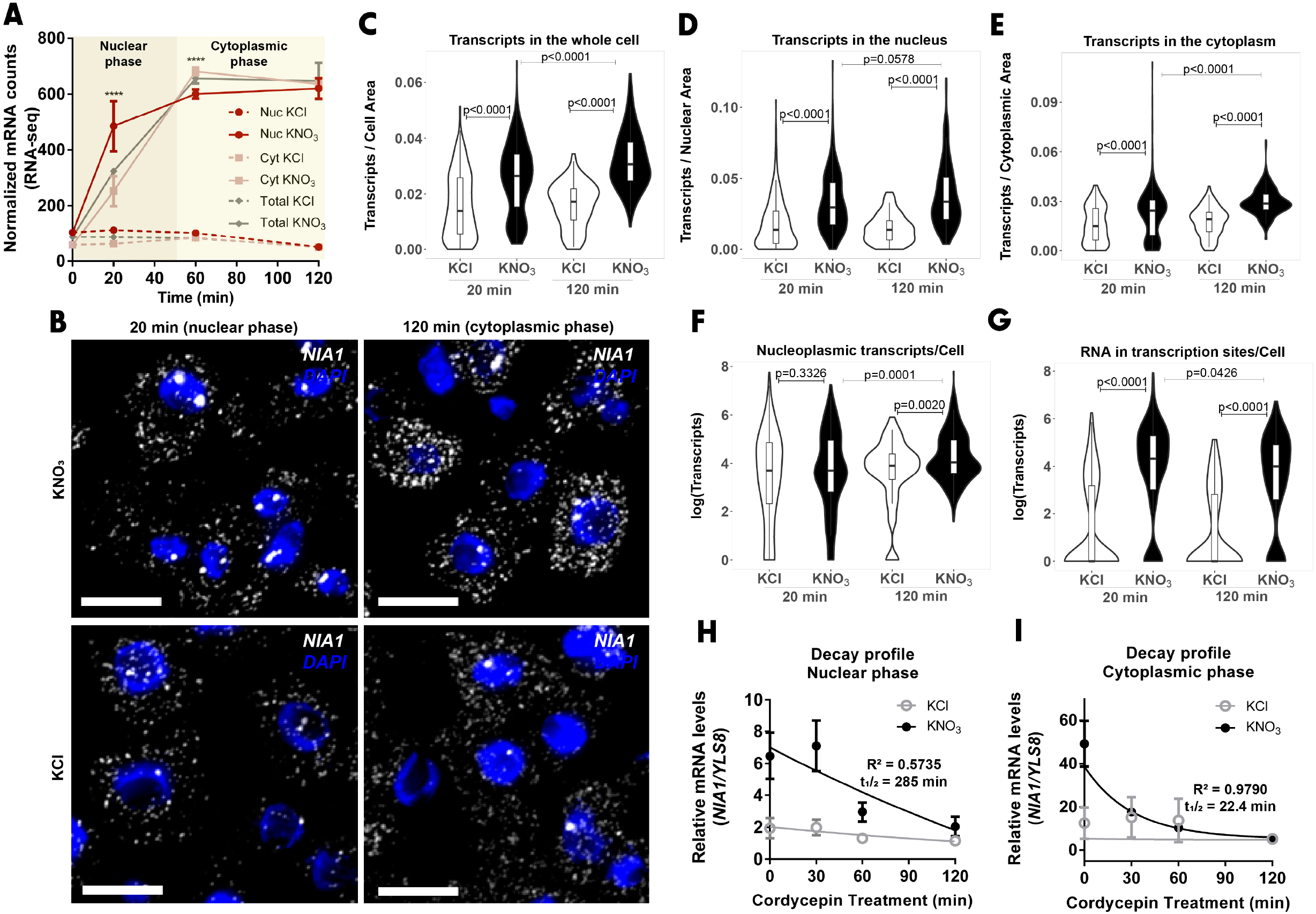
RNA single-molecule FISH detection and decay profiles for *NIA1* transcript in nuclear and cytoplasmic accumulation phases. **(A)** mRNA levels in cellular fractions measured by RNA-seq. (****) indicates statistical differences between nuclear and cytoplasmic fractions. **(B)** Representative microscopy deconvoluted images for *NIA1 in situ* detection in Arabidopsis root cells by RNA single-molecule FISH (smFISH). White color corresponds to signal detected for *NIA1* specific fluorescent probes. The blue color corresponds to the DAPI stain. Scale bar = 10 µm. **(C-G)** Quantification of the RNA smFISH. Violin plots show the distribution for transcript quantification in the nitrate (KNO_3_, black) or control (KCl, white) conditions at 20 or 120 min after the treatment. Boxes inside show the interquartile range (IQR – 25-75%), indicating the median value as a horizontal line. Whiskers show the ±1.58xIQR value. **(C)** Estimated number of transcripts per cell area in whole cells (nucleus+cytoplasm). **(D)** Number of transcripts per nuclear area. The number of nucleoplasmic transcripts and the estimated number of transcripts in transcription sites are included. **(E)** Number of cytoplasmic transcripts per area. **(F)** Number of active transcription sites per cell. **(G)** Estimated number of transcripts in each transcription site. **(H-I)** Comparison of *NIA1* half-lives (t_1/2_) after 20 min (nuclear phase) or 120 min (cytoplasmic phase) of KNO_3_ or KCl treatments. RNA levels were determined by RT-qPCR. Half-lives and coefficients of determination for regression models are indicated in each graph.

Furthermore, to evaluate the relationship between decay and the nuclear or cytoplasmic accumulation phases, we measured RNA levels by RT-qPCR after cordycepin treatments and compared differences in *NIA1* decay rates at 20 min and 120 min of nitrate treatments (Figure 6H-I). In its nuclear-accumulation phase, the *NIA1* transcript showed a half-life 12.7 times greater than in its cytoplasmic-accumulation phase. We obtained similar results for *VRN1*, another gene with a similar localization profile (Supplemental Figure 12A), that showed a 2.11-fold higher half-life at 20 min compared to 120 min (Supplemental Figure 12B-C).

These results provide evidence for a relationship between differential subcellular localization and mRNA stability for these nitrate-responsive genes. These results also suggest a role for nucleocytoplasmic dynamics in controlling transcript levels of rapidly replaced mRNAs. In the case of delayed-cytoplasmic transcripts (particularly *NIA1*), the control of the mRNA release from the DNA locus, and subsequent nuclear-to-cytoplasm export, could explain the lag in cytoplasmic mRNA accumulation. This strategy could avoid large quantities of newly synthesized mRNA (in the first minutes after the nitrate’s perception) overwhelming translation machinery. It could also play a role in coordinating the expression of multiple genes required for specific biological processes to operate in response to nitrate.

## DISCUSSION

This study described genome-wide mRNA nucleocytoplasmic dynamics in response to nitrate treatment in *Arabidopsis thaliana* roots. In addition to identifying new nitrate-responsive genes, we described 402 differentially localized transcripts (DLTs) in response to nitrate treatment using nuclear and cytoplasmic transcriptome data. These DLTs have relevant functions in the nitrate response and have characteristic sequence features associated with their localization pattern. Most of the induced DLTs showed big changes in synthesis and decay rates in response to nitrate treatments, indicating they are rapidly turned over. Our results suggest that controlling mRNA nucleocytoplasmic distribution is a strategy to fine-tune gene expression for mRNAs transcribed in bursts of gene expression. These findings highlight the relevance of modulating mRNA localization for controlling gene expression during the plant’s adaptive response to nitrogen nutrient signals.

### mRNA nucleocytoplasmic dynamics is regulated for relevant genes in the plant’s nitrate response

Using cell-fractionation and RNA-seq analysis, we obtained a high-resolution subcellular transcriptome in response to nitrate treatment in *Arabidopsis thaliana* roots. Thousands of genes have been previously reported as differentially expressed in response to nitrate treatments under various experimental conditions (Wang et al., 2004; Krouk et al., 2010; Canales et al., 2014; Varala et al., 2018; Alvarez et al., 2019; Swift et al., 2020), and several gene expression layers have been described (Vidal et al., 2020; Alvarez et al., 2021). Notwithstanding, we identified 1,183 regulated genes in the subcellular fractions that are not detected as regulated in the total fraction. A large proportion of these genes were not detected as nitrate-regulated in previous studies. This result indicates that our approach provides new information about mRNA accumulation in response to nitrate treatments, describing the mRNA levels in subcellular compartments and identifying new genes that have not been characterized in the *Arabidopsis thaliana*’s response to nitrate.

The nitrate response is a dynamic process. For instance, transcripts associated with nitrogen uptake and assimilation are enriched among regulated genes at early time points (5-15 minutes). Furthermore, other metabolic and developmental processes are regulated later (after the first hour) (Varala et al., 2018). Our work described the temporal dynamic of mRNA accumulation in subcellular fractions and showed that transcripts with different functions accumulate at different time points in the nucleus or cytoplasm. Furthermore, we identified a group of genes with delayed-cytoplasmic accumulation, which accumulates in the nucleus at early time points and later in the cytoplasm. These results indicate that transcript localization is fast and dynamically regulated in response to nitrate treatments. Two previous studies reported a similar temporal fraction-dependent regulation in plants when the nuclear transcriptome was compared with the cellular fraction (total poly(A)+) in response to hypoxia conditions (Lee and Bailey-Serres, 2019; Reynoso et al., 2019). These studies identified nuclear-retained transcripts under hypoxia associated with other stress functions. The authors proposed this mechanism as a strategy for minimizing the energetic demands after conditions of limited reserves in different plant species (Lee and Bailey-Serres, 2019; Reynoso et al., 2019), highlighting the plants’ ability to change transcripts’ availability in response to stimuli in a fast and dynamic manner.

Unlike previous evidence for nucleocytoplasmic mRNA localization in plants (Lee and Bailey-Serres, 2019; Reynoso et al., 2019), our study is not focused on a stressful condition, but it also represents a stimulus where energy optimization is crucial for plant growth and development (Vidal et al., 2020). Our results provide functional evidence for mRNA nucleocytoplasmic dynamics regulating gene expression. On the one hand, and comparable to previous reports in yeasts and plants under stress conditions (Saavedra et al., 1996; Zander et al., 2016; Lee and Bailey-Serres, 2019; Yeap et al., 2019), we observed that mRNAs with critical biological functions (e.g., nitrate metabolism) are enriched in the cytoplasm, indicating their accumulation is favored to sustain translation of genes that play essential roles in response to the nutrient. On the other hand, we also identified enriched GO-terms associated with nitrogen processes (for example, nucleotide biosynthesis and nitrate transporters) in nuclear-accumulated transcripts. Considering that nuclear-retained transcripts diminish their association to polysomes and thus protein synthesis (Pastro et al., 2017; Benoit Bouvrette et al., 2018; Lee and Bailey-Serres, 2019; Reynoso et al., 2019), it is not clear why these transcripts are nuclear-retained. A possible explanation for this phenomenon is that those transcripts could increase their levels in the cytoplasm after the 120 minutes analyzed, similar to the delayed-cytoplasmic accumulation pattern. The expression of these genes may not be required in rapid responses to the stimulus but for later processes, similar to that observed for general-function mRNAs in yeasts in response to heat stress (Saavedra et al., 1996; Zander et al., 2016). Alternatively, the accumulation of some transcripts in response to nitrate treatments may be an unintended consequence of activating a chromatin domain (Zhao et al., 2009; De and Babu, 2010). Therefore, nucleocytoplasmic regulation is relevant to fine-tune gene expression and coordinate the plant response to nitrate treatments.

### RNA-binding proteins may control RNA localization in response to nitrate treatments

Possible mechanisms explaining localization patterns are associated with variations in sequence features (Palazzo and Lee, 2018). In agreement with previous work in flies, humans, and parasites (Solnestam et al., 2012; Pastro et al., 2017; Benoit Bouvrette et al., 2018), we observed that cytoplasmic-accumulated mRNAs are shorter and have lower GC content than nuclear-accumulated transcripts. We also observed that mRNAs that accumulate in the nucleus are longer and have higher splicing-junction density than those enriched in the cytoplasm, agreeing with studies in animals and protozoa (Pastro et al., 2017; Benoit Bouvrette et al., 2018). A higher frequency of splicing sites could increase the processing time or the probability for intron retention, inhibiting their release from the DNA locus and delaying their cytoplasmic delivery (Custódio et al., 1999; Pandya-Jones et al., 2013; Ietswaart et al., 2017; Monteuuis et al., 2019) Sequence features are directly connected with RNA-binding protein (RBP) specificity. Particular nucleotide sequence motifs and secondary structures are determinants for protein-RNA recognition (Silverman et al., 2013; Gosai et al., 2015; Dedow and Bailey-Serres, 2019). Transcripts that share biological functions work as RNA regulons, coordinating their splicing, export, translation, and degradation (Keene, 2007; Culjkovic-Kraljacic and Borden, 2018). A positive correlation between nuclear accumulation, mRNA length, and specific motif-containing RBPs was described in Drosophila and human cells (Benoit Bouvrette et al., 2018). Furthermore, mRNAs with cytoplasmic enrichment have lower free energy for the predicted secondary structure in carcinoma cells (Solnestam et al., 2012). Our data showed differences among DLTs in the predicted RNA folding energy, suggesting transcripts with different localization patterns could bind to different nitrate-regulated RBPs.

Plants have a higher diversity of RBPs than other eukaryotic species, suggesting a better ability to adapt stimuli responses (Marondedze et al., 2016; Köster et al., 2017; Dedow and Bailey-Serres, 2019). For instance, the *A. thaliana* genome encodes four orthologs for the human export protein RBP ALYREF (ALY1, ALY2, ALY3, and ALY4) (Pfaff et al., 2018). Nevertheless, these RBPs have specific functions, considering that only ALY1 is associated with RNA-directed DNA-methylation transcripts in inflorescences (Choudury et al., 2019). Our analysis identified 131 out of 426 differentially expressed genes with ‘mRNA binding protein’ annotated molecular function in response to nitrate treatments. These results suggest that specific RBPs could promote RNA regulon formation to control functionally related mRNAs localization in response to nitrate treatments, as observed in other eukaryotic species.

### Differential RNA localization tunes gene expression for transcripts with rapid turnover

Our evidence supports a role for RNA synthesis and degradation in modulating nuclear and cytoplasmic mRNA levels, considering the synthesis occurs in the nucleus and most of the degradation in the cytoplasm (Łabno et al., 2016). A previous study in plants had proposed that nuclear accumulation of transcript is related to the activation of RNA synthesis when they observed higher transcript levels for nuclear-enriched RNA in Arabidopsis embryos (Palovaara and Weijers, 2019). Our data lend support to this hypothesis. DLTs with nuclear enrichment patterns have significant increments in RNPII occupancy and high transcript accumulation in synthesis loci by smFISH in response to nitrate treatments. Our data indicate that nuclear-accumulated transcripts are mostly newly synthesized after the nitrate treatment and do not reach the cytoplasm at the same rate they are transcribed, generating a differential nucleocytoplasmic distribution.

In agreement with our results, a negative correlation for RNA synthesis and decay has been reported for yeast, mouse, and fly cells under basal conditions (Miller et al., 2011; Tippmann et al., 2012; Chen and Van Steensel, 2017). Interestingly we found a stronger negative correlation when only DLTs were considered, comparable to the observed in *Saccharomyces cerevisiae*, where transcripts with specific functions in the osmotic stress response showed a stronger negative correlation (Miller et al., 2011). The increase of all RNA kinetic rates is a strategy for controlling transient induction and diminishing transcriptional noise (Rabani et al., 2014). For instance, rapid turnover occurs co-translationally in plants during the response to excess-light stress for a faster tuning of the genetic response to the stimulus (Crisp et al., 2017). This evidence suggests that DLTs undergo a faster replacement, probably due to their specific role in the cellular response to nitrate. Control of mRNA nucleocytoplasmic distribution would be another regulatory layer contributing to the expression of these transcripts.

### RNA nuclear export as a mechanism for buffering cytoplasmic transcript levels in response to nitrate treatments

Modulating mRNA nuclear export rates is a strategy for controlling transient cellular responses in different cellular and environmental stimuli. For example, mRNAs with stage-specific functions change between nuclear and cytoplasmic compartmentation during *Trypanosoma cruzi* development, regulating their expression (Pastro et al., 2017). Furthermore, mRNA nuclear export is regulated in transcripts with stress response functions in Drosophila (Chen and Van Steensel, 2017) and Arabidopsis (Lee and Bailey-Serres, 2019). Export, synthesis, and cytoplasmic decay rates are sufficient to predict nucleocytoplasmic mRNA levels using mathematical models (Bahar Halpern et al., 2015; Battich et al., 2015; Hansen et al., 2018), indicating that the contribution of other outputs for mRNA levels such as mRNA nuclear degradation (Das et al., 2003) and extracellular export (Thieme et al., 2015) cannot be considered in some cases. Our results show that nucleocytoplasmic accumulation of transcripts is a dynamic process whereby hundreds of genes change their distribution between cellular compartments in response to nitrate treatments. The synthesis and decay features for DLT transcripts do not explain the differential distribution in the nucleus and cytoplasm by themselves, suggesting a role for mRNA nuclear export modulation.

This potential nuclear export control is evident for the delayed-accumulated transcripts, which showed a temporal decoupling of nuclear and cytoplasmic mRNA-level increment. As we previously described, transcripts from the delayed-cytoplasmic accumulation pattern have a strong synthesis in response to nitrate. A low export rate would be required to maintain these high steady-state levels in the nucleus. In contrast, a high export rate would be required to keep cytoplasmic levels high during later times when the synthesis rate. Besides, delayed-cytoplasmic accumulated transcripts showed the highest destabilization among DLT patterns, increasing decay rates when the transcripts are more accumulated in the cytoplasm. This evidence suggests a connection between decay and export. A positive correlation between decay and export has been previously reported in flies (Chen and Van Steensel, 2017) and provides evidence on the importance of controlling the export rate to maintain cytoplasmic mRNA levels under cell requirements.

In this context, we propose mRNA nuclear export as a mechanism for buffering the expression levels of critical genes for the nitrate response. The temporal retention of mRNAs in the nucleus is a strategy for controlling the expression of transcripts synthesized during bursts of transcription (Bahar Halpern et al., 2015; Tudek et al., 2019). This can be achieved, for instance, by blocking the release of the transcripts from the gene locus and avoiding them from reaching the nuclear pore complex (Katahira et al., 2019; Singh et al., 2019). Our results are consistent with this model. In particular, *NIA1* showed a delayed-cytoplasmic accumulation in response to nitrate, with early nuclear retention at transcription sites. This gene encodes an enzyme involved in the first step of nitrate reduction (Cheng et al., 1988; Santos-Filho et al., 2014). This is a committed step that is subject to multiple levels of regulation (Yanagisawa, 2014; Krapp, 2015), including transcriptional regulation (Zhao et al., 2018) and mRNA decay (Wu et al., 2020). A recent study showed that *NIA1* degradation generates many siRNAs, some of which regulate their own expression, allowing the plant to quickly adapt its metabolism to the nutritional state (Wu et al., 2020). Thus, regulating *NIA1* mRNA localization could allow postponing the turnover of this transcript to efficiently regulate the plant nutritional status in response to nitrate treatments.

We described the mRNA nucleocytoplasmic dynamics in response to nitrate in Arabidopsis roots. The patterns observed for differentially localized transcripts can be partially explained by characteristic mRNA sequence features, synthesis, or decay rates. We propose that controlling nuclear-to-cytoplasm delivery is a strategy for buffering RNA levels and fine-tune gene expression for transcripts that undergo a fast turnover. Understanding how nitrate regulates the expression of genes involved in metabolism, growth, and development is essential for developing new biotechnological solutions in agriculture. Our research gives new insights into plants’ post-transcriptional RNA regulation and provides the basis for elucidating the role of mRNA nuclear export in the context of nitrogen nutrition.

## METHODS

### Plant growth and nitrate treatments

*Arabidopsis thaliana* seedlings (Col-0 ecotype) were grown in hydroponic media for 15 days, using ammonium succinate as the only nitrogen source in the Phytatray^TM^ system (Sigma, Cat.P1552). Treatments with KNO_3_ (or KCl as control) to a final concentration of 5 mM were performed, according to Alvarez et al., 2014). Root tissue was collected at 0, 20, 60, and 120 min of treatment and immediately frozen in liquid nitrogen until processing.

### RNA extraction from cellular fractions

Cell subfractionation was achieved through differential centrifugation in a sucrose solution according to the protocol published by Xu and Copeland, (2012): the pellets obtained correspond to the nuclear fraction, and the supernatants collected correspond to the cytoplasmic fraction. Unfractionated tissue was stored from ground roots for ‘total’ RNA extraction. RNA extraction from all cellular fractions was performed using an acid phenol-chloroform protocol published by Darnell, (2012). Finally, all extracted RNA samples were purified following the Clean-up for Liquid Samples protocol from PureLink® RNA Mini Kit (Ambion, Cat, 12183018A). Concentration, integrity, and purity parameters were evaluated for RNA extractions by capillary electrophoresis (Fragment Analyzer, STANDARD SENSITIVITY RNA ANALYSIS KIT DNF-471, Advanced Analytical Technologies) and spectrophotometry (Nanodrop2000, Thermo Scientific), procuring to have more than two micrograms of RNA, RNA Quality Number (RQN) higher than 6.0, and optimal absorbance ratios (A260/A280 and A260/A230) for each extraction.

### RT-qPCR measurements

cDNA was synthesized from nuclear and cytoplasmic RNA using Improm II RT (Promega, Cat. #A3800), and cDNA levels were measured by qPCR using the Brilliant III Ultra-Fast qPCR Kit (Agilent Technologies, Cat. #600880) and the StepOnePlus^TM^ qPCR System (Agilent Technologies). Primers listed in Supplemental Table 2 were used for qPCR measurements. cDNA levels were calculated using the LinRegPCR software (Ramakers et al., 2004).

Nuclear transcript enrichment was evaluated by RT-qPCR detection of specific regions of unprocessed transcripts in genes with constitutive expression [*CLATHRIN COAT ASSEMBLY*, *RAN3*, and *EIF4G*]. Primers were designed to detect intronic regions for unprocessed RNAs and between two exons (at the end of an exon and the beginning of the closest neighbor exon) for processed RNAs (Supplemental Table 2). The analysis was performed from cDNA synthesized from RNA extractions obtained for different cellular fractions using random primers (Promega, Cat. #C1181). Unprocessed RNA levels were normalized using the mean value of three processed mRNAs.

Differential RNA levels in the cellular fractions were confirmed, measuring *MPK9*, *SDR2, SUFE2*, *RCAR1, NRT2.2*, *BCA4, BZIP3*, *AT1G49230*, *NIA1,* and *IDH1* (representative transcripts for DLT localization patterns). The mean RNA levels for *CLATHRIN COAT ASSEMBLY* and *PP2AA3* were used as a normalizer factor. For RNA decay evaluation by qPCR, *NIA1* and *VRN1* levels were measured from cordycepin-treated plants, using the mean level of *RAN3* and *MON1* as a normalizer factor. For both experimental designs, the best pair of normalizer genes was validated following the strategy described by Remans et al., (2014), using NormFinder (Andersen et al., 2004). cDNA was synthesized using the oligo(dT) 5’-TTTTTTTTTTTTTTTTTTTTV-3’.

### RNA sequencing from cellular fractions

cDNA libraries (from PolyA enriched RNA) were prepared by Macrogen service (South Korea), using TruSeq® Stranded mRNA LT Sample Preparation Kit (Illumina, Cat. RS-122-2101). Libraries were synthesized with RNA from each cellular fraction (nuclear, cytoplasmic, and total) for control (KCl) and treated (KNO_3_) conditions for the four time-points collected (0, 20, 60, and 120 minutes). Libraries were sequenced in the Illumina Novaseq6000 platform with 100 bp paired-end reads by Macrogen.

### RNA-seq data analysis

R software software packages (CRAN R Project) were used for most data analysis. The FastQC software (0.10.0 version, Babraham Bioinformatics) was used to check the reads’ quality, and then the sequences were processed with Trimmomatic v0.36 (Bolger et al., 2014) for removing the low-quality reads. The sequences were mapped to the *Arabidopsis thaliana* genome (Araport11 annotation) using HISAT2 (Kim et al., 2015), and finally, Rsubread R Library (Liao et al., 2013) was used for calculating the number of reads in Transcripts Per Million (TPM).

We used quantile normalization to identify differentially expressed genes in the cellular fractions (Smyth, 2005). This strategy was considered the best for reducing the bias generated by the different conditions of the subcellular fractions. Nuclear and cytoplasmic TPMs were quantile normalized together (for comparisons between cellular fractions), and total TPMs were analyzed separately. To identify genes that are differentially accumulated by the treatment and change during the time-course, a two-way ANOVA model was performed from the quantile normalized counts (in log2 scale) of each cellular fraction, evaluating the effects of treatment (KCl and KNO_3_), time (20, 60, 120 min) and their interaction through the model. In this way, transcripts that fit the model with a significant p-value for treatment (T) or its interaction with time (Treatment:Time) were considered as genes whose mRNA levels change within the cellular fraction in response to nitrate.

Given that nuclear and cytoplasmic fractions are not independent of each other, a ΔNC value (Normalized counts in nuclear fraction minus normalized counts in cytoplasmic fraction) was calculated to identify genes whose transcripts show different distributions between these cellular fractions. From these values, a similar analysis (as mentioned above) was performed. The transcripts whose ΔNC values fit the 2-way ANOVA model with a significant p-value (<0.01) for treatment or its interaction with time were considered differentially localized transcripts (DLTs) in response to the nitrate treatments.

The different lists of regulated genes were compared using the Sungear software (Poultney et al., 2007). The Multiple Experiment Viewer (MeV) software (Saeed et al., 2003) was used to visualize and cluster the data. Gene groups were defined by hierarchical clustering from their Pearson correlation, using an average linkage method and defining a threshold distance of 0.5. Enrichment analysis of Gene Ontology (GO) and Kyoto Encyclopedia of Genes and Genomes (KEGG) terms were performed using the BioMaps software from VirtualPlant v1.3 (Katari et al., 2010), selecting terms with a p-value with FDR (False Discovery Rate) correction lower than 0.05. GO terms were summarized with REVIGO (http://revigo.irb.hr) set to obtain a medium-size list of terms according to Resnik similarity (Supek et al., 2011). The sequence features of these transcripts were analyzed, extracting the information from the Generic Feature Format (GFF) file of Araport11 annotation for each gene’s most abundant isoform according to the RNA-seq data. Prediction of mRNAs secondary structure was performed using the RNAfold function from ViennaRNA Package 2.0 (Lorenz et al., 2011).

### RNA stability evaluation

*Arabidopsis thaliana* seedlings were treated with a transcription inhibitor after the treatment with the nutrient, and RNA decay rates and half-lives were calculated for each condition (KNO_3_ and KCl). 15-day old *Arabidopsis thaliana* seedlings were treated as described above. After 20 or 120 min of nutrient treatment, the plants were transferred to a solution of cordycepin 0.6 mM (Sigma Cat. #C3394) prepared in MS without nitrogen (PhytoTechnology Laboratory Cat. #M407) in a growth chamber with low agitation. Roots were collected at 0, 30, 60, and 120 min after the cordycepin treatment. RNA was extracted using TRIzol^TM^ reagent (Invitrogen Cat. 15596) following the protocol described by Macrae (2007). RNA was used for cDNA synthesis and subsequent quantification by RNA-seq (for RNA from seedlings treated for 120 min with nitrate) or qPCR to evaluate stability at other treatment times.

For RNA-seq analysis, twenty-four different libraries were synthesized from RNA extracted from three experiments (separate plant material grown independently). cDNA libraries (from rRNA-depleted RNA) were prepared by Macrogen service (South Korea) with TruSeq® Stranded Total RNA Library Prep Plant (Illumina, Cat. 20020611) using RNA from nitrate or control conditions after cordycepin treatment. Libraries were sequenced by Macrogen in Illumina Novaseq6000 platform with 100 bp paired-end reads, requesting 40 million reads per sample. Raw data were analyzed as described above for RNA-seq from cellular fractions until obtaining TPM normalized counts. The Multiple Experiment Viewer (MeV) (Saeed et al., 2003) software was used to visualize and cluster the data. Gene clusters were defined by hierarchical clustering from their Pearson squared correlation, using a complete linkage method, defining a threshold distance of 0.5.

Decay rates (k_decay_) and then half-lives (t_1/2_) were calculated by adjusting the measured RNA levels (C) as an exponential function of time (t). The mathematical adjustment for C(t) was developed assuming a constant decay rate, according to the function: C(t) = e^-kdecay * t^ (Gutierrez et al., 2002; Narsai et al., 2007; Sorenson et al., 2018). ‘RNA decay’ R-package (Sorenson et al., 2018) was used for decay modeling for RNA-seq data. Models in which decay rate changed or not between KNO_3_ and KCl treatments were evaluated, and the model with the lowest Akaike Information Criterion (AIC) statistics was selected.

### RNA polymerase II occupancy changes

Data from RNA Polymerase II Chromatin Immunoprecipitation sequencing (RNPII-ChIPseq) was analyzed (Alvarez et al., 2019). This data was obtained from *Arabidopsis thaliana* roots treated for 12 min with nitrate in the same conditions used to identify DLTs. Normalized sequence counts in regions between 500 bp upstream the TSS and 500 bp downstream the TTS were evaluated by differential accumulation with DESeq2 package (Love et al., 2014), calculating a fold change of RNPII occupancy between treated (KNO_3_) and control condition (KCl).

### RNA single-molecule FISH

RNA single-molecule Fluorescence *in situ* Hybridization (RNA smFISH) was performed according to the protocol described in Duncan et al., (2017). Forty-eight probes were designed using the Stellaris Probe Designer software (version 2.0 from Biosearch Technologies) to recognize exonic regions for the *NIA1* transcript. Probes with Quasar670 fluorophores were synthesized by Stellaris. Probe sequences are listed in Supplemental Table 3.

Fifteen-day-old *A. thaliana* seedlings were treated with nitrate (and KCl as control) for 20 and 120 min. Some of these plants were also treated with cordycepin 0.6 mM for 120 min after nutrient treatment for transcription site analysis. Roots were collected, fixed in 4% paraformaldehyde solution, and squashed on microscope slides to obtain cell monolayers. Fixed samples were hybridized with the probe set and then with DAPI 100 ng/mL. The visualization and imaging were performed with a Zeiss LSM800 inverted microscope, using an x63 oil-immersion objective and a cooled quad-port CCD (charge-coupled device) ZEISS Axiocam 503 mono camera. The following wavelengths were used for fluorescence detection: for Quasar670, an excitation filter 625-655 nm was used, with signal detection at 665-715 nm; for DAPI, an excitation filter of 335-383 nm with signal detection at 420–470 nm. For all experiments, a series of optical sections with z-steps of 0.22 µm were collected. Maximum projections and analysis of three-dimensional pictures were performed using Fiji. For image deconvolution and quantification, FISH-quant software was used (Mueller et al., 2013). Tutorial instructions for batch analysis for "Mature mRNA quantification" and "Nascent mRNA quantification" were followed (Mueller et al., 2013).

### Accession numbers

Accession numbers based on The Arabidopsis Information Resource (TAIR) (https://www.arabidopsis.org) for all genes examined in this study are: *NIA1* (AT1G77760), *RAN3* (AT5G55190), *EIF4G* (AT3G60240), *CLATHRIN COAT ASSEMBLY PROTEIN* (AT4G24550), *PP2AA3* (AT1G13320), *MON1* (AT2G28390), *MPK9* (AT3G18040), *SDR2* (AT3G51680), *SUFE2* (AT1G67810), *RCAR1* (AT1G01360), *NRT2.2* (AT1G08100), *BCA4* (AT1G70410), *BZIP3* (AT5G15830), *IDH1* (AT4G35260), *VRN1* (AT3G18990). Sequence data from this article can be found in the National Center for Biotechnology Information Gene Expression Omnibus under the project accessions: PRJNA720236 (data from cellular fractions) and PRJNA791353 (data from stability assays).

## Supplemental Data files

### Supplemental Data Sets

**Supplemental Data Set 1.** Differentially expressed genes in response to nitrate treatments in subcellular fractions.

**Supplemental Data Set 2.** Over-represented GO-terms for differentially expressed genes in response to nitrate treatment in cellular fractions.

**Supplemental Data Set 3.** List of DLTs in response to nitrate treatments.

**Supplemental Data Set 4.** Over-represented GO-terms in DLTs.

**Supplemental Data Set 5.** Over-represented KEGG-terms in DLTs.

**Supplemental Data Set 6.** RNA polymerase II occupancy changes for DLTs.

**Supplemental Data Set 7.** Decay profiles and estimated half-lives for DLTs.

**Supplemental Data Set 8.** Ranking for RNA polymerase II occupancy and half-life changes for DLTs.

## ACKNOWLEDGMENTS

We thank Luis Villarroel, Ph.D. (Pontificia Universidad Católica de Chile) for help with data normalization and statistics strategy. This work was supported by grants from the Fondo de Desarrollo de Áreas Prioritarias (FONDAP), Center for Genome Regulation (15090007), and the Agencia Nacional de Investigación y Desarrollo (ANID), through ANID–Millennium Science Initiative Program-Millennium Institute for Integrative Biology (iBio), FONDECYT 1180759 to R.A.G and Ph.D. scholarship (21161516) to A.F.

## AUTHOR CONTRIBUTIONS

A.F. and R.A.G. designed this research. A.F. wrote the manuscript, with corrections from T.M., S.R., and R.A.G. A.F., T.M, and R.A.G designed the RNA-seq data analysis pipeline. A.F. performed all the experiments that involved bench work and analyzed the data. T.M. analyzed the RNAseq data. S.R. designed, supervised, and discussed smFISH experiments. R.A.G supervised the study.

## COMPETING INTERESTS

The authors declare no competing interests.

## SUPPLEMENTAL MATERIAL

**Supplemental Figure 1.** Unprocessed transcripts are enriched in nuclear fractions and reduced in cytoplasmic fractions.

RNA levels measured by RT-qPCR for unprocessed transcripts in the different cellular fractions (Total, nuclear, and cytoplasmic). RNA was extracted from root tissue from nitrate- or control-treated seedlings for 60 min (KNO_3_ or KCl, respectively). Three different constitutive-expressed transcripts were detected: **(A)** *EIF4G*, **(B)** *CLATHRIN COAT ASSEMBLY PROTEIN*, and **(C)** *RAN3*. RNA values were normalized with mean processed RNA levels. Detection was performed using primers flanking (processed RNA) or inside (unprocessed RNA) intronic regions for each gene. Bars represent the mean ± standard error of 3 independent experiments. Statistical analysis was performed using ANOVA/Tukey’s HSD test. Different letters above bars denote statistically significant differences (ANOVA/Tukey’s HSD test, p<0.05).

**Supplemental Figure 2.** Number of differentially expressed genes in cellular fractions

The lists of genes with significant factors obtained by two-way ANOVA analyses of RNAseq data is represented using the Sungear tool (Poultney et al., 2007). The triangle shows the factors at the vertexes (Treatment, time, and the interaction between both). The circles inside the triangle represent the genes controlled by the different factors, as indicated by the arrows around the circles. The size of each circle is proportional to the number of genes associated with that circle. The number of genes in the circle is shown next to the corresponding circle. **(A)** Total fraction, **(B)** Nuclear fraction, **(C)** Cytoplasmic fraction, and **(D)** Nuclear-Cytoplasmic subtraction. Differentially expressed genes in response to nitrate treatments are colored in red.

**Supplemental Figure 3.** Differentially expressed genes in response to nitrate treatment in cellular fractions.

**(A)** Sungear representation for comparing the lists of differentially expressed genes in Total, nuclear, and cytoplasmic fractions (vertexes) in response to nitrate (KNO_3_) compared with control (KCl) treatments. Circles indicate the number of genes shared among each list.

**(B)** Heatmap with mRNA levels in cellular fractions for differentially expressed genes in subcellular fractions that are not identified in the total fraction. Genes were hierarchically clustered using the correlation of mRNA levels in the nuclear and cytoplasmic fractions (panel I). mRNA levels in the total fraction are shown in panel II. Each column represents the mRNA levels for one replicate under each condition. Nuclear (Nuc) or Cytoplasmic (Cyt) regulation is indicated on the figure’s right side. Three replicates from independent experiments are shown as separated columns for each condition.

**Supplemental Figure 4.** Comparison of regulated genes in response to nitrate treatments from different transcriptomic studies

The lists of differentially expressed genes in response to nitrate in different transcriptomic studies are represented using the Sungear tool (Poultney et al., 2007). Vertexes of the polygon show different lists for ALVAREZ_2019 (Alvarez et al., 2019), CANALES_2014 (Canales et al., 2014), KROUK_2010 (Krouk et al., 2010), Swift_2019 (Swift et al., 2020), VARALA.ROOT_2018 (Results for root tissue in Varala et al., 2018), WANG.ROOT_2004 (Results for root tissue in Wang et a., 2004), TOTAL_FRACTION, NUCLEAR_FRACTION, and CYTOPLASMIC_FRACTION (results from this work). The circles inside the polygon represent the identified genes from each work, as indicated by the arrows around the circles. The size of each circle is proportional to the number of genes associated with that circle. Genes only identified in cellular fractions are colored in red, and the number of these genes in each list is indicated as text.

**Supplemental Figure 5.** DLTs in response to nitrate treatments that are not regulated in the total fraction.

**(A)** Sungear representation for comparing lists of total differentially regulated genes and DLTs in response to nitrate treatment.

**(B)** Heatmap with mRNA levels in cellular fractions for DLTs that are not identified in the total fraction. Genes were hierarchically clustered using the correlation of mRNA levels in the nuclear and cytoplasmic fractions (panel I). mRNA levels in the total fraction are shown in panel II. Each column represents the mRNA levels for one replicate under each condition.

**Supplemental Figure 6.** DLT expression clusters for the different localization patterns in response to nitrate treatments.

We identified 13 expression clusters (Figure 1) for DLTs in response to nitrate treatments. The localization pattern (NR – Nuclear reduction, CR – Cytoplasmic reduction, NA – Nuclear accumulation, D – Delayed-cytoplasmic accumulation, and CA – Cytoplasmic accumulation) each cluster represents is indicated. DLTs in response to nitrate treatments were separated by the cellular fraction where the main changes are observed (Nuclear or cytoplasmic) and if they show an accumulation or reduction in the nitrate condition. Graphs show mean values of z-scored normalized RNA levels (orange line) and 95% confidence interval for mean values for three biological replicates (shadow).

**Supplemental Figure 7.** qPCR validation for differential accumulation of representative DLTs.

mRNA levels for representative genes from DLT localization patterns in response to nitrate. The mRNA levels were analyzed at the time-point, where the most significant differences between nuclear and cytoplasmic levels were observed in RNAseq data. The left panels show the mRNA levels in all time-courses measured by RNA-seq. The asterisks indicate statistical differences between nuclear and cytoplasmic levels in a specific time-point (Two-way ANOVA, Tukey post-test. *; p<0.05; **: p<0.01, ***: p<0.001, ****: p<0.0001). The right panels show the mRNA levels measured by RT-qPCR. The different letters denote statistically significant differences (Two-way ANOVA, Tukey post-test, p<0.05).

**(A)** RNA levels for *MPK9* (nuclear reduction – NR - pattern). qPCR measurements were performed at 20 min of treatment.

**(B)** RNA levels for *SDR2* (nuclear reduction – NR - pattern). qPCR measurements were performed at 120 min of treatment.

**(C)** RNA levels for *SUFE2* (cytoplasmic reduction – CR - pattern). qPCR measurements were performed at 120 min of treatment.

**(D)** RNA levels for *RCAR1* (cytoplasmic reduction – CR - pattern). qPCR measurements were performed at 120 min of treatment.

**(E)** RNA levels for *NRT2.2* (nuclear accumulation – NA - pattern). qPCR measurements were performed at 120 min of treatment.

**(F)** RNA levels for *RCA4* (nuclear accumulation – NA - pattern). qPCR measurements were performed at 120 min of treatment.

**(G)** RNA levels for *BZIP3* (cytoplasmic accumulation – CA - pattern). qPCR measurements were performed at 20 min of treatment.

**(H)** RNA levels for *AT1G49230* (cytoplasmic accumulation – CA - pattern). qPCR measurements were performed at 20 min of treatment.

**(I)** RNA levels for *NIA1* (delayed cytoplasmic accumulation – D - pattern). qPCR measurements were performed at 20 and 120 min of treatment.

**(J)** RNA levels for *IDH1* (delayed cytoplasmic accumulation – D - pattern). qPCR measurements were performed at 20 and 120 min of treatment.

**Supplemental Figure 8.** Extended sequence-related features for DLTs in response to nitrate treatments.

Sequence features of the most abundant isoform for nitrate-regulated genes, according to Araport11 annotation. Violin plots show the distribution of **(A-C)** length features, **(D-E)** GC content, and **(G-H)** free energy for optimal secondary structure prediction (RNAfold software) for DLTs localization patterns (NR – Nuclear reduction, CR – Cytoplasmic reduction, NA – Nuclear accumulation, D – Delayed-cytoplasmic accumulation, and CA – Cytoplasmic accumulation). We also include nitrate-regulated genes in the total fraction that are not differentially localized as a control (TA – Total accumulated, or TR – Total reduced). Boxes inside show the interquartile range (IQR – 25-75%), indicating the median value as a horizontal line. Whiskers show the ±1.58xIQR value. We compared the distributions using one-way ANOVA and Dunnett post-test. We include p-values and brackets to highlight relevant comparisons.

**Supplemental Figure 9.** Sequence features for DLTs in response to nitrate treatments separated by expression clusters.

Sequence features of the most abundant isoform for DLTs, according to Araport11 annotation. Box plots show minimum to maximum (bars) and 5-95 percentile (boxes) distributions. Clusters were grouped based on their localization pattern patterns (NR – Nuclear reduction, CR – Cytoplasmic reduction, NA – Nuclear accumulation, D – Delayed- cytoplasmic accumulation, and CA – Cytoplasmic accumulation). The mean value from all DLTs is indicated in the dashed line. (*) show statistical differences (p<0.05) among clusters and DLTs distribution from One-way ANOVA, Tukey post-test analysis. **(A-D)** Length features **(E-H)** GC content features **(I-K)** Free energy for optimal secondary structure prediction (RNA fold software) **(L)** Splicing junction density (Calculated as two times the number of introns divided by exonic region length). **(M)** Changes in RNPII occupancy after 12 min of treatment. Values are graphed as the fold change (FC) of the KNO_3_/KCl ratio in a logarithmic scale (log2). **(N)** Changes in half-life after 120 min of treatment. Values are graphed as the fold change (FC) of the KNO_3_/KCl ratio in a logarithmic scale (log2).

**Supplemental Figure 10.** Decay profiles for DLTs in response to nitrate treatments.

Decay profiles for the 402 DLTs in response to nitrate treatments. RNA levels were measured by RNAseq from KNO_3_ (Left panel) or KCl (Right panel) treated roots for 120 min and then treated with cordycepin for 0, 30, 60, or 120 min. The RNA levels were normalized using the mean value at 0 min (T0) of cordycepin treatment. Genes were hierarchically clustered in six groups according to their decay profiles in the KNO_3_ condition (indicated in the left part of the figure). The localization patterns with which each DLT belongs are indicated in the right part of the figure. Replicates from independent experiments are shown as separate columns for each condition.

**Supplemental Figure 11.** Transcription sites analysis by RNA single-molecule FISH during nitrate treatment.

**(A)** Representative microscopy deconvoluted images from two independent experiments for *NIA1* detection in *Arabidopsis* root cells from nitrate (KNO_3_, top) or control (KCl, bottom) for 20 min, and then treated with cordycepin 0.6 mM for 0 (left) or 120 (right) min. The white signal corresponds to specific probes for *NIA1* associated with Quasar670 fluorophore. The blue signal corresponds to the DAPI stain. Scale bar = 10 µm.

**(B-C)** Transcription site analysis. Violin plots show the distribution for transcript quantification in the nitrate (KNO_3_, black) or control (KCl, white) condition after 20 or 120 min of treatment (Figure 6E). Boxes inside show the interquartile range (IQR – 25-75%), indicating the median value as a horizontal line. Whiskers show the ±1.58xIQR value. One-way ANOVA Tukey post-test p-values for significant differences are indicated (n.s: non-significant, p-value>0.1). Four images from different roots from two independent experiments were quantified for each time/condition. **(B)** Number of active transcription sites per cell. **(C)** Estimated number of transcripts by each active transcription site.

**Supplemental Figure 12.** Decay profiles for *VRN1* transcript in response to nitrate treatments

**(A)** mRNA levels in cellular fractions measured by RNA-seq for *VRN1.* Asterisks (****) indicate statistical differences between nuclear and cytoplasmic fractions, according to two-way ANOVA, Tukey post-test. SEM from three independent experiments is shown as error bars.

**(B-C)** Comparison of half-lives (t_1/2_) for *VRN1* between nuclear **(B)** and cytoplasmic **(C)** phases. RNA levels were determined by RT-qPCR. Measured half-lives and the determination coefficient for linear regression are indicated for each graph. SEM from three independent experiments is shown as error bars.

## SUPPLEMENTAL TABLES

**Supplemental Table 1.** RNA-seq libraries parameters from total and cellular fractions.

| Parameter | Total | Nuclear | Cytoplasmic |
| --- | --- | --- | --- |
| Library concentration range (nM) | 22.49 - 356.94 | 72.53 - 136.50 | 50.03 - 119.51 |
| Library size range (bp) | 316 - 353 | 274 - 300 | 270 - 292 |
| Number of reads (in millions) [Mean ± SD] | 68.92 ± 9.26 | 70.57 ± 7.33 | 70.75 ± 5.37 |
| Q<sub>30</sub> (%) [Mean ± SD] | 94.72 ± 0.31 | 93.98 ± 0.19 | 93.69 ± 0.32 |
| Reads after quality filters (%) [Mean ± SD] | 93.31 ± 0.54 | 89.56 ± 0.44 | 88.92 ± 0.60 |
| Mapped reads to the genome (%) [Mean ± SD] | 92.01% ± 1.44% | 86.05% ± 3.80% | 84.12% ± 6.91% |
| Correlation between replicates [Mean ± SD] | 0.988 ± 0.002 | 0.984 ± 0.003 | 0.984 ± 0.003 |

**Supplemental Table 2.**
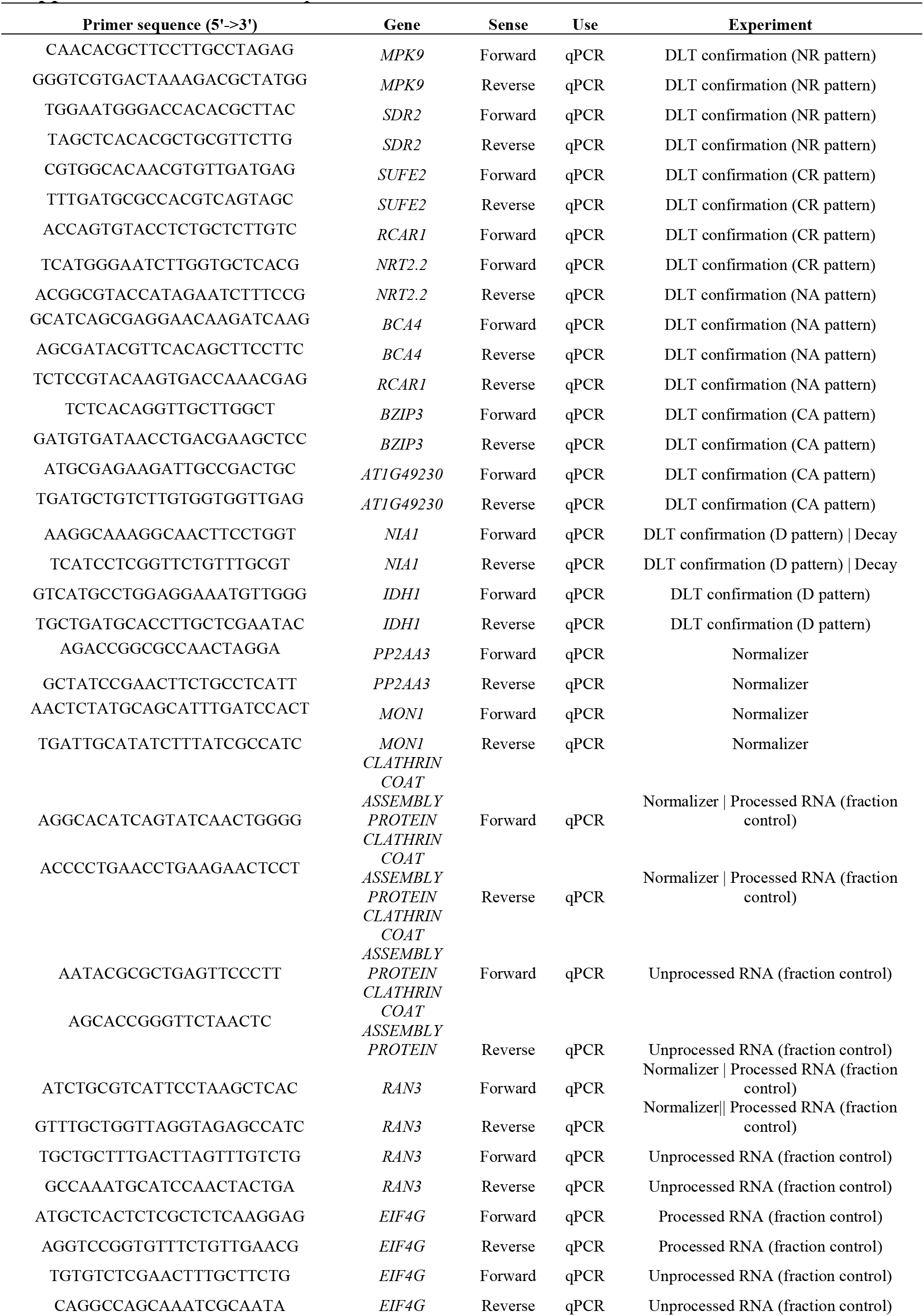
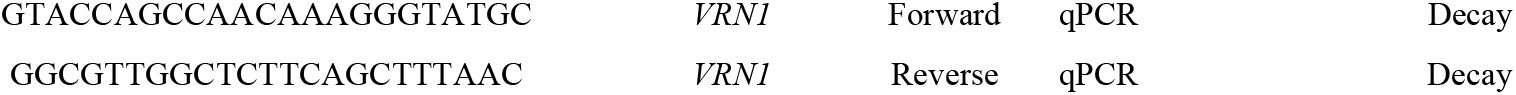
List of primers

**Supplemental Table 3.**
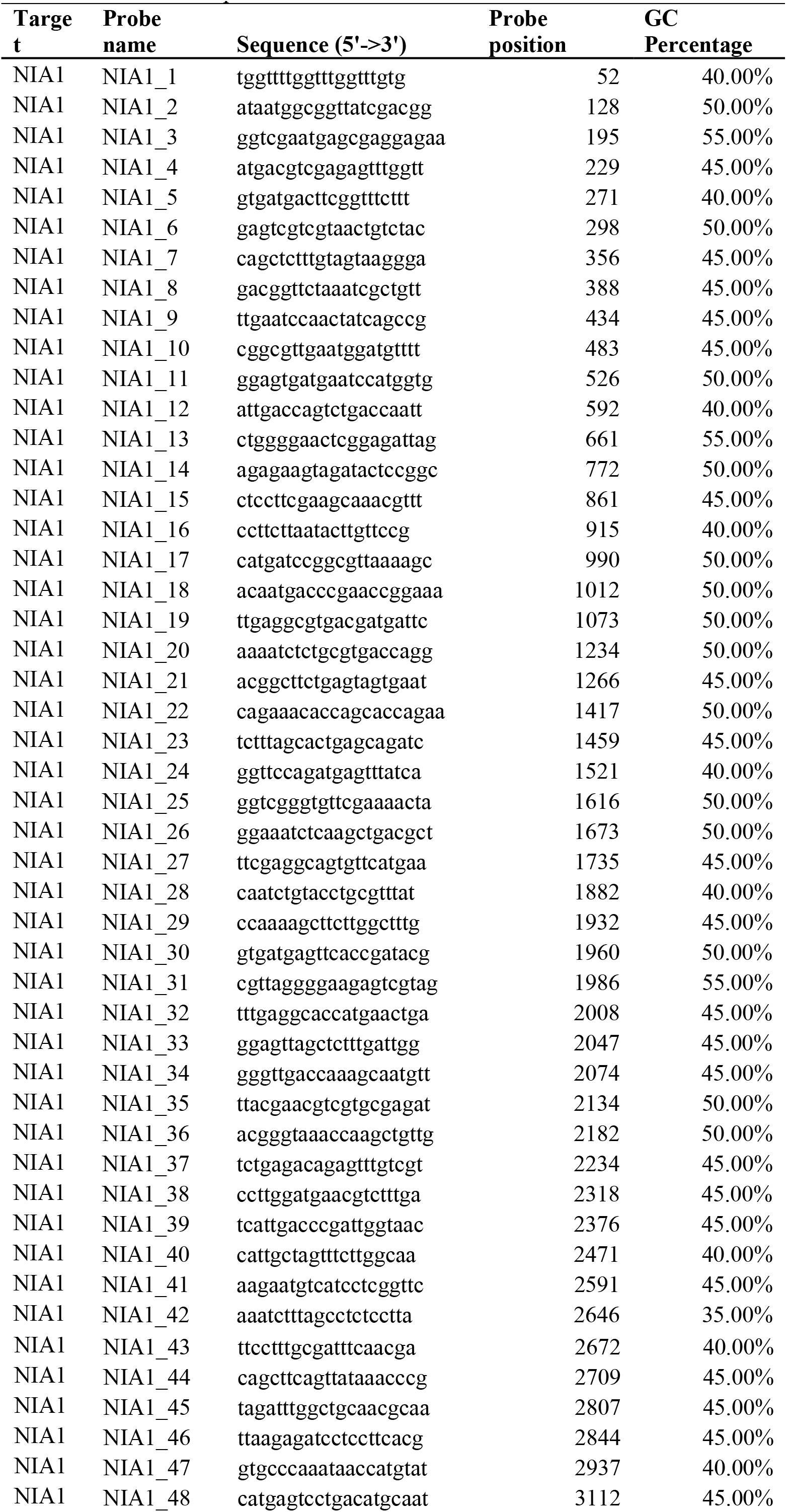
List of probes for *NIA1* RNA smFISH

